# Dominance patterns in dynamic environments

**DOI:** 10.1101/2020.02.25.965103

**Authors:** Archana Devi, Kavita Jain

## Abstract

Natural environments are seldom static and therefore it is important to ask how a population adapts in a changing environment. We consider a finite, diploid population with intermediate dominance evolving in a periodically changing environment and study how the fixation probability of a rare mutant depends on its dominance coefficient and the rate of environmental change. We find that in slowly changing environments, the dominance patterns are the same as in the static environment, that is, if a mutant is beneficial (deleterious) when it arrives, it is more (less) likely to fix if it is dominant. But in fast changing environments, these patterns depend on the mutant’s fitness on arrival as well as that in the time-averaged environment. We find that in a rapidly varying environment that is neutral or deleterious on-average, an initially beneficial (deleterious) mutant that arises while selection is decreasing (increasing) has a fixation probability lower (higher) than that for a neutral mutant leading to a reversal in the standard dominance patterns. We also find that recurrent mutations decrease the phase lag between the environment and the allele frequency, irrespective of the level of dominance.

## 1 Introduction

Natural environments change with time and a population must continually adapt to keep up with the varying environment (Gillespie, 1991; Messer *et al*., 2016; Bleuven and Landry, 2016). It is therefore important to understand the adaptation dynamics of a finite population subject to both stochastic and environmental fluctuations. This is, in general, a hard problem but some understanding of such dynamics has been obtained in previous investigations. For example, when the environment changes very rapidly, on the time scale of a generation, the adaptation dynamics are simply determined by the time-averaged environment (Gillespie, 1991).

Environments can, of course, vary slowly and recent experiments have shown the impact of rate of change in the environment on the population fitness (Salignon *et al*., 2018; Boyer and Sherlock, 2019). The fixation probability, fixation time and adaptation rate in changing environments have also been studied in a number of theoretical work (Takahata *et al*., 1975; Gillespie, 1993; Mustonen and Lässig, 2008; Assaf *et al*., 2008; Uecker and Hermisson, 2011; Waxman, 2011; Peischl and Kirkpartick, 2012; Cvijović et al., 2015; Dean et al., 2017), and it has been found that when the environment changes at a finite rate, the population dynamics are strongly determined by the environment in which the mutant arose. In particular, the dependence of fixation probability on arrival times of the mutant has been demonstrated numerically for very large populations by Uecker and Hermisson (2011), but the analogue of their result for a finite population is not known.

All the works mentioned above assume a haploid or a diploid population with semi-dominance (Charlesworth, 1998; Bourguet, 1999). In a static environment, the fixation probability of a dominant beneficial mutant is known to be higher than when it is recessive (*Haldane’s sieve*) (Haldane, 1927), while the opposite pattern holds if the mutant is deleterious (Kimura, 1957). How these results are affected in dynamic environments is, however, not known.

In this article, we therefore study the adaptation dynamics of a finite, diploid population with intermediate dominance evolving in an environment that changes periodically due to, for example, seasonal changes or drug cycling. We find that in dynamic environments, the magnitude of the fixation probability of a rare mutant differs substantially from the corresponding results in the time-averaged environment. Furthermore, the dominance patterns in the fixation probability can differ from those expected in the static environment depending on the rate of environmental change, the arrival time of the mutant and its fitness in the time-averaged environment. However, when recurrent mutations occur, our results for the average allele frequency and population fitness suggest that dominance does not have a strong influence.

## 2 Model

We consider a finite, sexually-reproducing diploid population of size *N* with a single biallelic locus under selection. The (Wrightian) fitness of the three genotypes denoted by *aa, aA, AA* is 1 + *s*, 1 + *hs*, 1, respectively, where the dominance coefficient 0 < *h* ≤ 1. The population evolves under a periodically changing environment that is modeled by a time-dependent selection coefficient 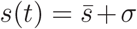 sin *ωt* where 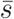 is the selection coefficient averaged over a period 2*π*/*ω*; in the following, we assume that 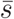 is arbitrary but *σ* > 0. We ignore random fluctuations in the environment so that selection changes in a predictable fashion. Mutations occur at a constant symmetric rate *μ* between the two alleles. Mating is assumed to be random and therefore the genotypic frequencies are in Hardy-Weinberg equilibrium at the time of conception; this simplification allows us to work with allele frequency *x* and 1 − *x* of the allele *a* and *A*, respectively (Ewens, 2004).

We study the model described above analytically using an appropriate perturbation theory and numerically through stochastic simulations that were carried out using a continuous time Moran model in which the number *i* of allele *a* increases and decreases by one at rate *r_b_* and *r_d_*, respectively, given by

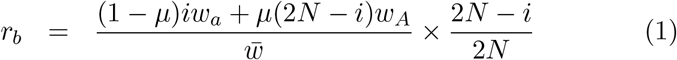

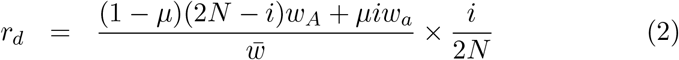

where

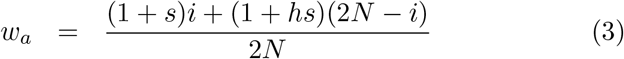

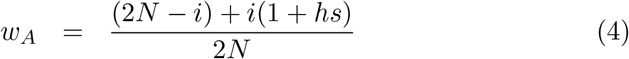

are the marginal fitness of the allele *a* and *A*, respectively, and 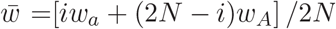 is the population-averaged fitness. We choose the time at which either birth or death happens from an exponential distribution with rate *r_b_*+*r_d_*, and then allow one of the events to occur with a probability proportional to its rate (also see, for e.g., Uecker and Hermisson (2011)).

Below we first consider the weak mutation regime (4*Nμ* ≪ 1) to understand in detail how the fixation probability of a rare mutant depends on various environmental and population factors such as the driving frequency and dominance parameter. We also briefly explore the strong mutation regime (4*Nμ* ≫ 1), and calculate the population fitness and the phase lag between the allele frequency and changing environment.

## 3 Fixation probability of a rare mutant

In a static environment, the fixation probability of a mutant allele can be described by a backward Kolmogorov equation (Ewens, 2004). For the model described in the last section, the probability *P*_fix_ that mutant allele *a* present in frequency *x* at time *t* fixes eventually is given by

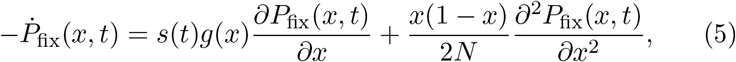

where dot denotes a derivative with respect to time and *g*(*x*) = *x*(1 − *x*)(*x*+*h*(1 − 2*x*)). On the right-hand side (RHS) of the above equation, the first term describes the deterministic rate of change in the allele frequency (Ewens, 2004) and the second term captures the stochastic fluctuations due to finite population size. Since the left-hand side (LHS) of the above equation is nonzero, (5) is *inhomogeneous* in time; that is, the eventual fixation probability depends on the arrival time *t_a_* (or phase *θ_a_* = *ωt_a_*) of the mutant (Uecker and Hermisson, 2011; Waxman, 2011).

Equation (5) does not appear to be exactly solvable, and approximate methods such as perturbation theory require an exact solution of the unperturbed problem (*σ* = 0) which is not known in a closed form for nonzero 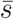 (Kimura, 1957). Therefore to obtain an analytical insight, we study the fixation probability using a branching process for positive 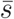 and analyze the above diffusion equation for 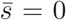 using a time-dependent perturbation theory. Some numerical results for negative 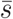 are given in Sec. S1. Before proceeding to a quantitative analyses, we first give a qualitative picture of the process in the following section.

### 3.1 Qualitative features

Figure 1 shows the fixation probability of a single mutant arising in an environment that is neutral on an average 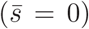. In a slowly changing environment, the dominance relationships are found to be the same as in the static environment (that is, a dominant mutant that starts out as a beneficial (deleterious) one has a higher (lower) chance of fixation). This is because at small frequencies, as Fig. 2a illustrates, fixation occurs rapidly so that the sign of selection remains the same from arrival to fixation.

**Figure 1:**
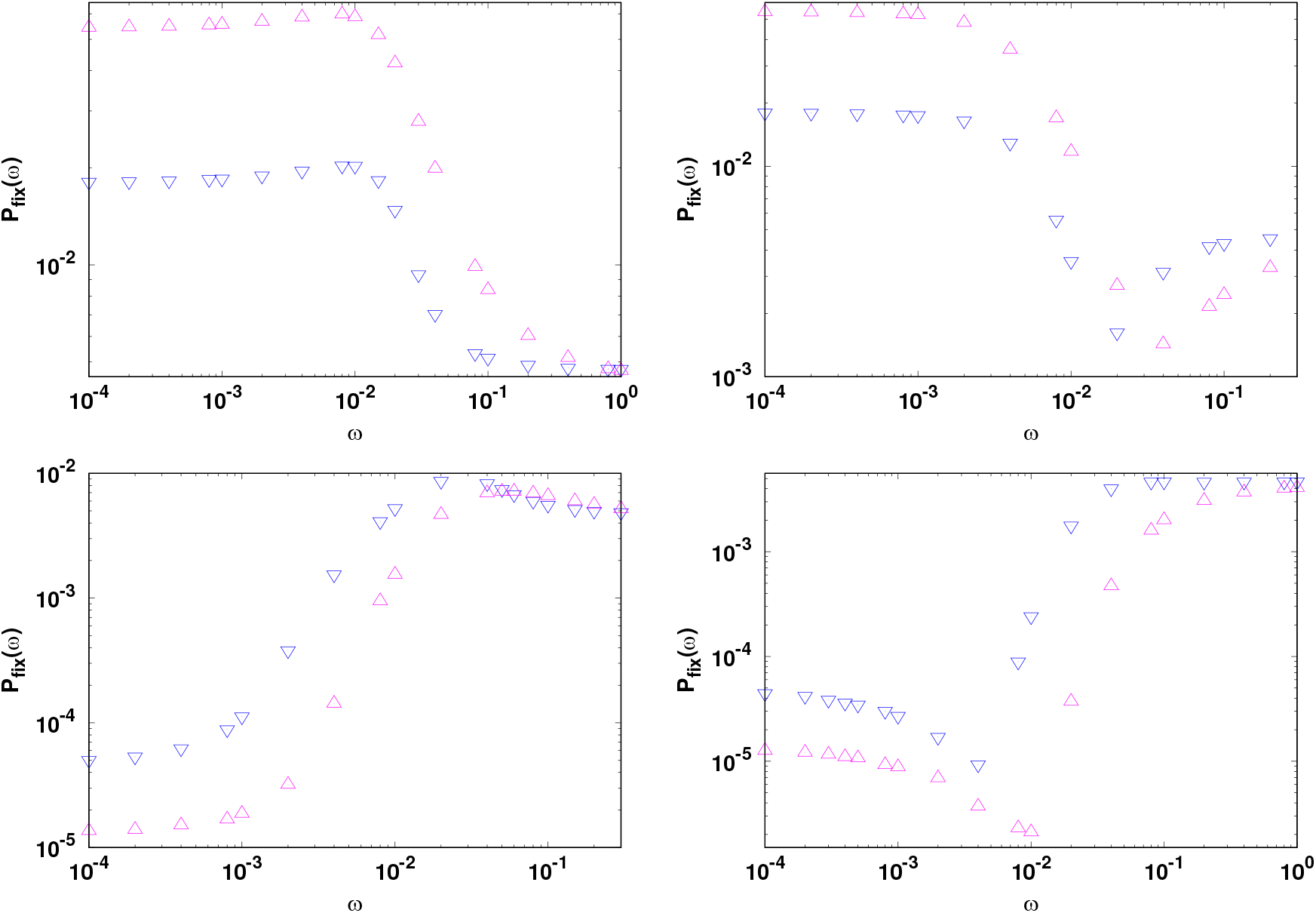
Fixation probability as a function of driving frequency for a mutation that is neutral on an average and arises in a large population of size *N* ≫ *σ*^−1^ for *h* = 0.1(∇) and 0.9(Δ), and *θ_a_* = *π*/4, 3*π*/4,5*π*/4, 7*π*/4 (clockwise from top left). The other parameters are *N* = 100 and *σ* = 0.1.

In rapidly changing environments, as Fig. 1 demonstrates, the dominance patterns can differ from those in static environments. For large frequencies, although the fixation times are much longer than the time period of environmental change (see Fig. 2b), the arrival time still plays an important role in determining the chance of fixation as the mutant must escape the stochastic loss at short times (in fact, Fig. 2b shows that the effect of genetic drift is strongest in the first seasonal cycle). Then a mutant that arises while selection is positive but the selection strength is decreasing will soon encounter an environment with negative fitness effects that affect a dominant mutant more adversely than the recessive one leading to a lower chance of fixation of the dominant mutant.

**Figure 2:**
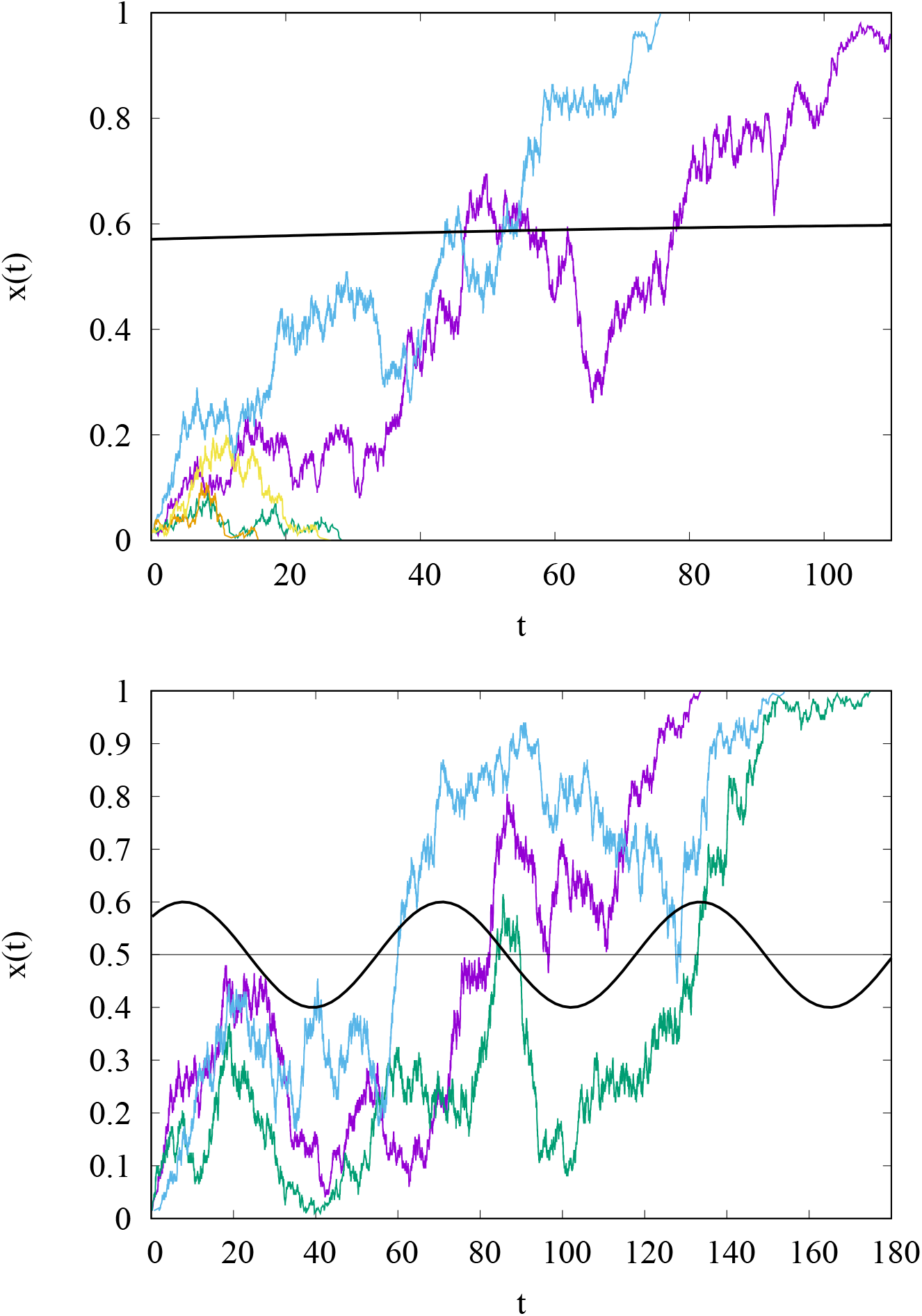
Allele frequency runs starting with a single mutant for *N* = 200, 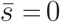, *σ* = 0.1, *h* = 0.5, *θ_a_* = *π*/4 for *ω* = 0.005 (top) and 0.1 (bottom). The solid line shows the selection coefficient *s*(*t*) shifted to show on the scale.

### 3.2 Beneficial mutant in a large population

When the mutant is beneficial on an average 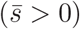 and the population is large enough 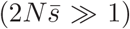, one can use a branching process to find the fixation probability of a rare mutant. This parameter regime has been studied for a haploid population by Uecker and Hermisson (2011); generalizing their results to diploids, we find that

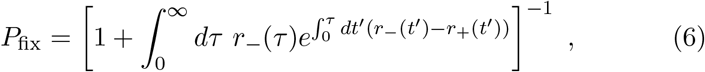

where *r*_−_(*t*) = lim_*N*→∞_*r_d_*/*i,r*_+_(*t*) = lim_*N*→∞_*r_b_/i*. From (1) and (2), we obtain *r*_−_ = 1, *r*_+_ = 1 + *hs*; using these in (6), we find that

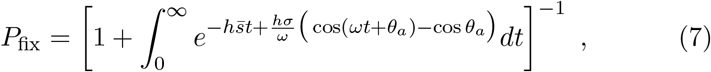

which reduces to (29) of Uecker and Hermisson (2011) for *h* = 1/2 and *ξ* = 1 (note that their expression contains a typographical error). The fixation probability (7) is, in general, a nonmonotonic function of driving frequency *ω* and strongly depends on the arrival phase *θ_a_* = *ωt_a_* (Uecker and Hermisson, 2011; Peischl and Kirkpartick, 2012).

Equation (7) is analyzed in Appendix A for 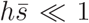. For 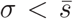 (and 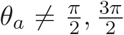) where the mutant is beneficial at all times, we find that

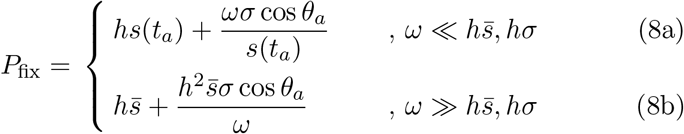

where 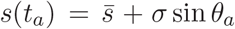. Since the fixation probability of a beneficial mutant with constant selection coefficient *s*_0_ is given by *hs*_0_ (Haldane, 1927), the above expressions show that for very slowly changing environment, the fixation probability is determined by the selection coefficient of the mutant at the instant it arose while for very rapidly changing environments, it depends on the time-averaged selection coefficient 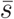 (Gillespie, 1993; Mustonen and Lässig, 2008).

The effect of slowly changing environment is captured by the deviation from *hs*(*t_a_*) in (8a) which changes linearly with the driving frequency and is *independent* of the dominance parameter. In contrast, for rapidly changing environments, the fixation probability (8b) is sensitive to dominance as the deviation from the asymptotic result 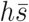 depends on *h* (also, see Fig. 3). The inset of Fig. 3 as also (8a) and (8b) show that the dominant mutant has higher fixation probability than the recessive one at *all* driving frequencies. In other words, Haldane’s sieve (Haldane, 1927) that favors the establishment of beneficial dominant mutations in static environments continues to operate in dynamic environments in which the mutant is beneficial at all times. Equations (8a) and (8b) also emphasize the important role of the arrival time of the mutant. If the beneficial mutant arises while the selection coefficient is increasing (decreasing) with time, the fixation probability at small frequencies increases (decreases) with *ω* and approaches the asymptotic value 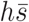 from above (below) at high frequencies.

**Figure 3:**
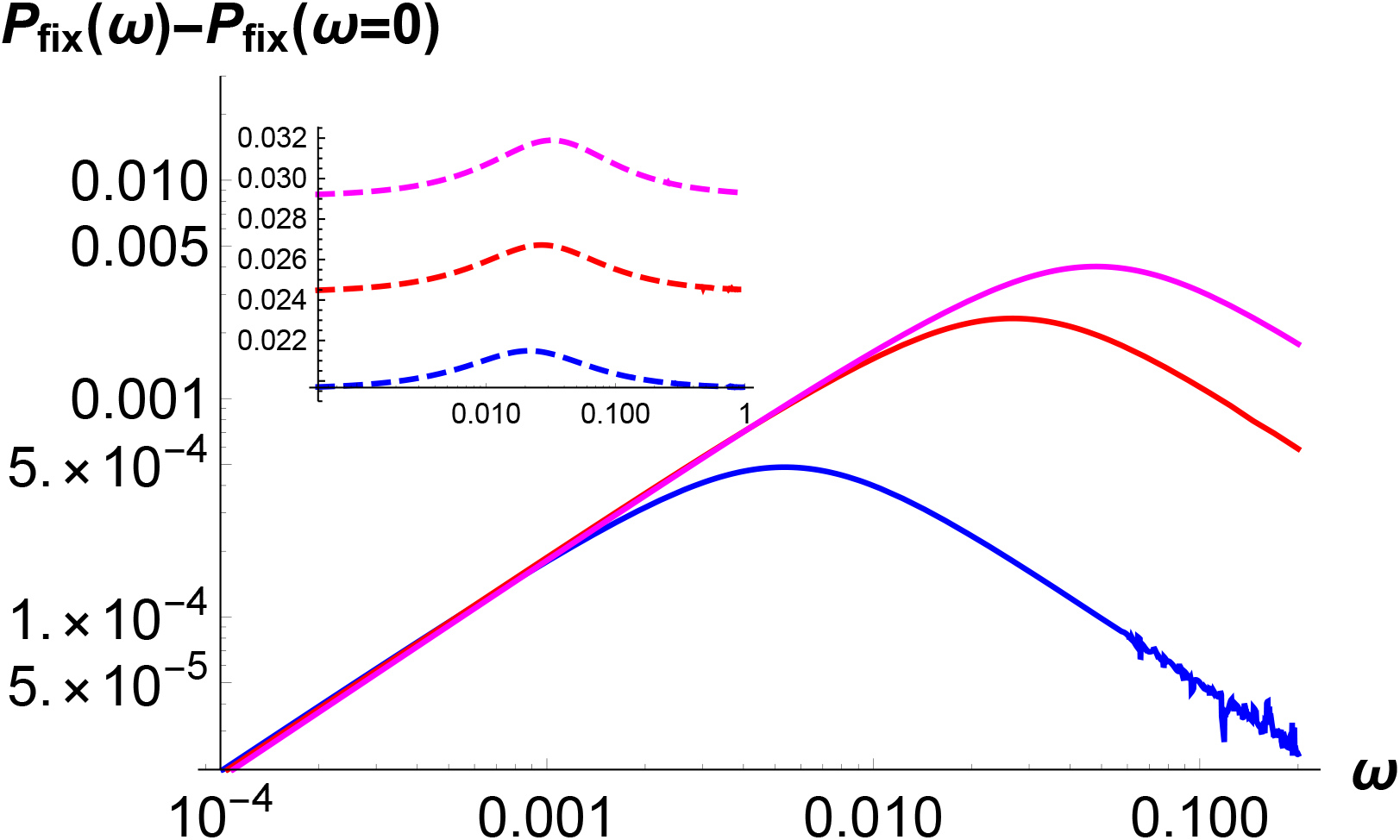
The inset shows the fixation probability *P*_fix_(*ω*) given by (7) for a mutant that is beneficial on an average for dominance coefficient *h* = 0.4, 0.5, 0.6 (bottom to top). In the main figure, the effect of changing environment is shown by subtracting the fixation probability *P_fix_*(*ω* = 0) = *hs*(*t_a_*)/[1 + *hs*(*t_a_*)] with 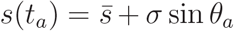 for *h* = 0.1, 0.5, 0.9 (bottom to top). In both plots, 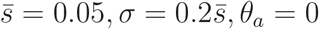.

On matching the expressions (8a) and (8b), the fixation probability is found to have an extremum at a *resonant frequency*,

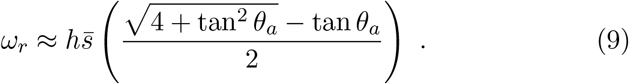

Thus a minimum or a maximum in the fixation probability occurs when the environment changes at a rate proportional to the ‘natural frequency’ of the population, *viz*., its average growth rate. As already mentioned above, this extremum is a minimum if the mutant arrives while selection is decreasing (*π*/2 < *θ_a_* < 3*π*/2) and a maximum otherwise. For the two special values, 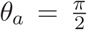 and 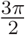, the fixation probability monotonically decreases and increases, respectively, with *ω*.

For 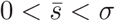, the mutant is not beneficial at all times; in this case, the expression (8a) for small frequencies holds if *s*(*t_a_*) > 0 (see Appendix A). Otherwise as the mutant is initially deleterious and arises in an infinitely large population, the fixation probability is essentially zero. However, (8b) for large frequencies is valid for any *s*(*t_a_*) as the time-averaged fitness is assumed to be positive. As shown in Fig. S2 for 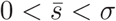, the dominant mutant has a higher fixation probability than the recessive one at large frequencies. But at small frequencies,the dominance pattern depends on whether the mutant is beneficial or deleterious on arrival.

### 3.3 Neutral mutant in a finite population

We now calculate the fixation probability of an on-average neutral mutant using the backward diffusion equation (5).

#### 3.3.1 Small population

We first consider a small population of size *N* ≪ *σ*^−1^ and analyze the *ω* ≪ *σ* and *ω* ≫ *σ* regimes in Appendix B and C, respectively. On using (B.8) and (C.2), we find that the fixation probability of allele *a* present in a single copy at time *t_a_* = *θ_a_/ω* is given by

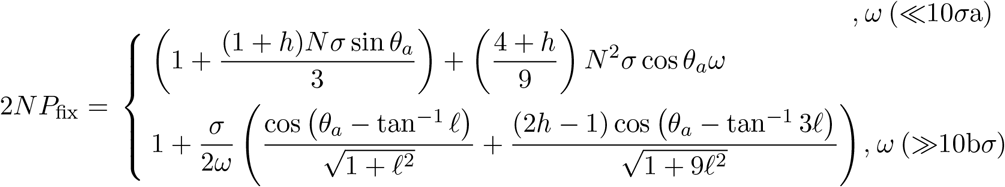

where *ℓ* = (*Nω*)^−1^. The expression (10b) holds for any *ω* ≫ *σ* but for *ω* ≫ *N*^−1^, it simplifies to

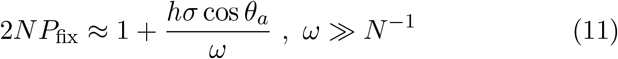

which monotonically approaches the fixation probability of a neutral mutant in a static environment. The above results show that the change in the fixation probability depends weakly on the dominance parameter when the environment changes slowly, but has a strong dependence on h in rapidly changing environments. The top panel of Fig. 4 shows that the expressions (10a) and (10b) are in very good agreement with the simulation results, and suggests that the resonant frequency does not depend on the dominance coefficient. As detailed in Appendix C, we find that

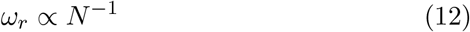

**Figure 4:**
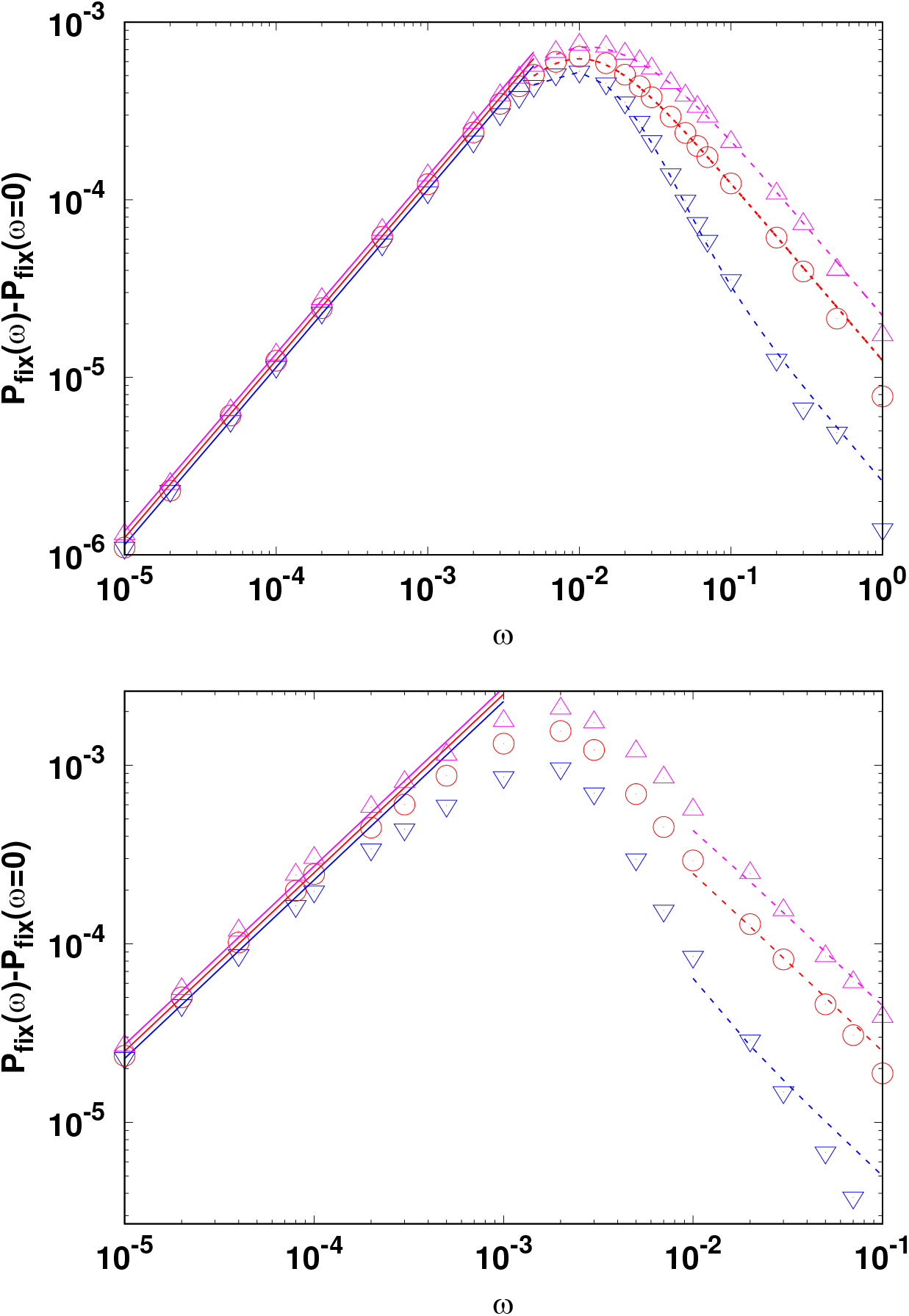
Effect of changing environment on the fixation probability of a mutant that is neutral on an average and arises in a small population (*N* ≪ *σ*^−1^, top) and large population (*N* ≫ *σ*^−1^, bottom). The points show the simulation data obtained by averaging over 10^7^ independent runs. The solid line shows the expression (10a) for small driving frequencies and the dashed lines represent (10b) for large frequencies. Here *N* = 100, *σ* = 0.005 (top) and *N* = 1000, *σ* = 0.01 (bottom), *θ_a_* = 0, and *h* = 0.1(∇), 0.5(°), 0.9(Δ). The value *P_fix_*(*ω* = 0) subtracted on the *y*-axis was obtained numerically.

and depends weakly on the dominance coefficient.

#### 3.3.2 Large population

We now consider large populations with size *N* ≫ *σ*^-1^. For *ω* ≪ *N*^−1^ ≪ *σ*, as discussed in Appendix B, we find that in slowly varying environments,

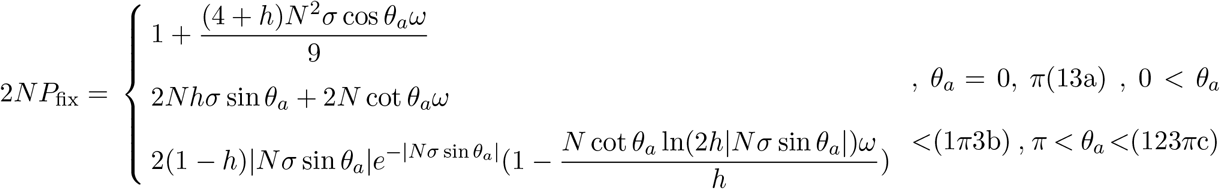

The above equations show that if the mutant arises when the selection is positive, the fixation probability increases linearly with *ω* if selection is increasing but decreases otherwise. The magnitude of the slope is, however, independent of *h* and *σ*. Similar statements apply for *s*(*t_a_*) ≤ 0. For large frequencies (*ω* ≫ *N*^−1^, *σ*), the fixation probability is given by (11) for any *s*(*t_a_*), and approaches the asymptotic neutral behavior from above (below) when *ṡ* = *ds/dt* is positive (negative) with the fixation probability increasing (decreasing) with increasing *h*. Figure 4 shows a comparison between our analytical and numerical results when the mutant arrives at *θ_a_* = 0 (for other arrival phases, see Fig. S3), and we find a good agreement.

Our perturbation expansions in Appendix B and C are not valid for intermediate frequencies (*N*^−1^ ≪ *ω* ≪ *σ*). However, our numerical simulations suggest that as for small populations, the resonant frequency scales as *N*^−1^ here also.

## 4 Average allele frequency and population fitness

We now turn to the strong mutation regime where 4*Nμ* ≫ 1 and briefly study the dynamics of the allele frequency in changing environments. For 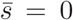, the frequency distribution Φ(*x*, *t*) of allele *a* under changing selection, mutation, and genetic drift obeys the following forward Kolmogorov equation (Ewens, 2004),

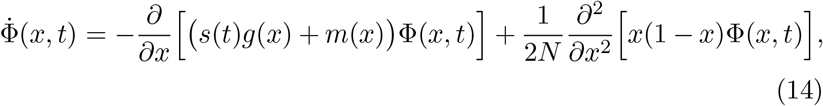

where the mutation term *m*(*x*) = *μ*(1 − 2*x*) and, as before, *s*(*t*) = *σ* sin*ωt*, *g*(*x*) = *x*(1−*x*)(*x*+*h*(1−2*x*)). The above equation is analyzed in Appendix D for small selection amplitude *σ*, and we find that at large times, the allele frequency distribution Φ(*x, t*) is given by (D.5).

To get an insight into how the allele frequency changes in changing environments, we find the population-averaged allele frequency,

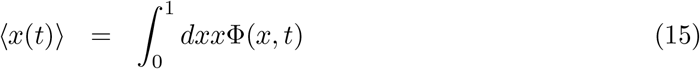

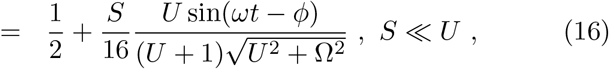

where *U* = 4*Nμ, S* = 4*Nσ*, Ω = 2*Nω*, 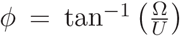. The above equation shows that 〈*x*(*t*)〉 oscillates about one half with the same driving frequency as *s*(*t*) but a different phase. The phase difference *ϕ* decreases with increasing mutation rate so that the allele frequency changes almost in-phase with the environment for *U* ≫ Ω but lags behind by a phase *π*/2 for *U* ≪ Ω. The latter behavior for rare mutations is already illustrated in Fig. 2 where the mutant’s allele frequency keeps increasing as long as the selection is positive and decreases when *s*(*t*) becomes negative. In contrast, for *U* ≫ Ω, the population keeps up with the environment as mutations occur faster than the time scale of environmental change.

Equation (16) also shows that the allele frequency amplitude remains close to the time-averaged amplitude when the environment changes rapidly. But it is significantly different from one half in slowly changing environments and varies nonmonotonically with the mutation rate with a maximum at the scaled mutation rate 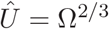. We also find that although the distribution Φ(*x, t*) depends on the dominance coefficient (see Appendix D), the average allele frequency (for small *σ*) is independent of *h*, as illustrated in the inset of Fig. 5.

**Figure 5:**
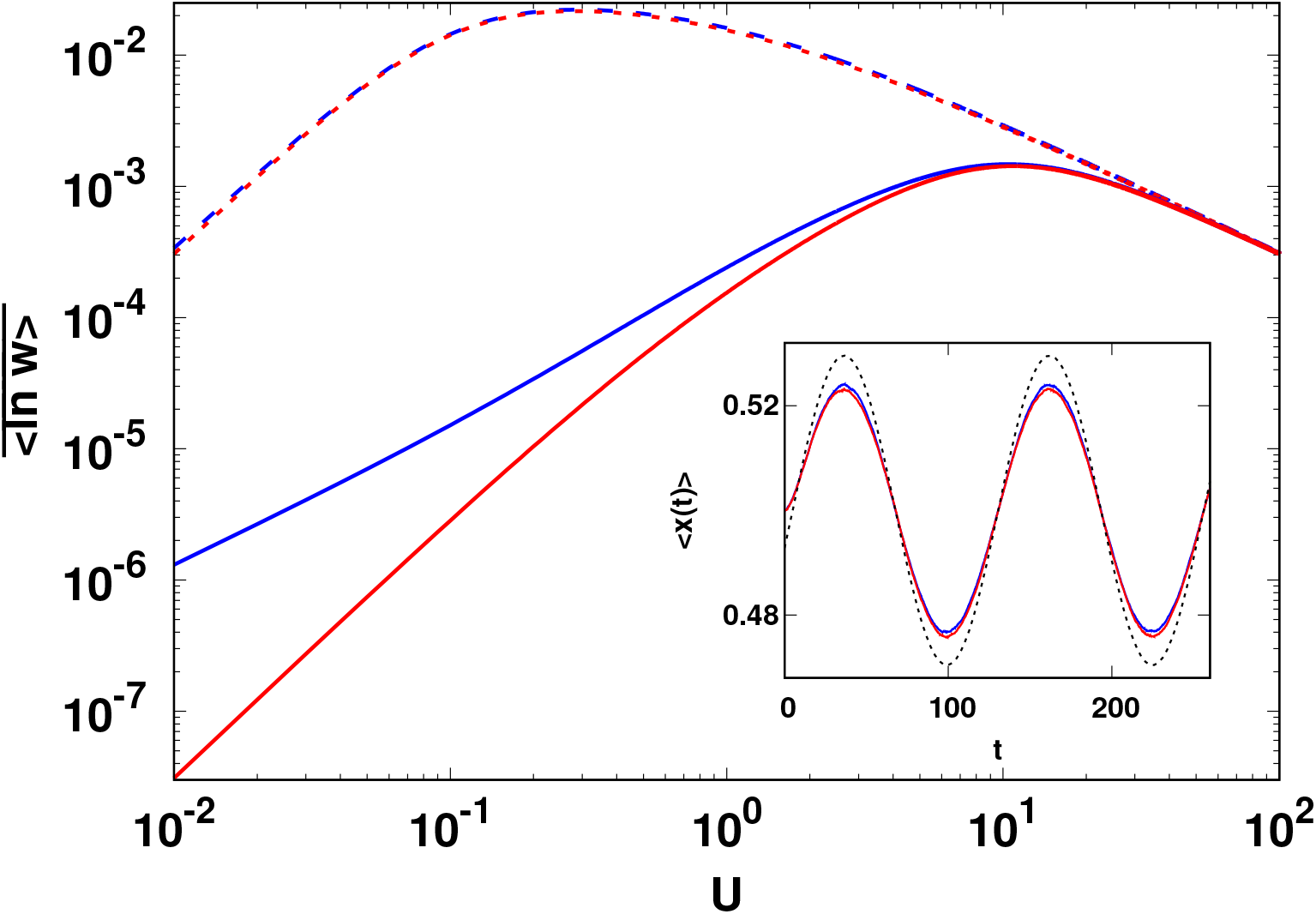
Main: Average fitness (18) in a periodically changing environment as a function of scaled mutation rate *U* = 4*Nμ* for scaled frequency Ω = 2*Nω* = 0.1 (dashed lines), 10 (solid lines). Inset: Dynamics of population-averaged allele frequency for *U* = 40 and Ω = 10. The black dotted line is the analytical formula (16) for the allele frequency. In all plots, *N* = 100, *σ* = 0.05 and *h* = 0.1 (blue) and 0.5 (red).

The above described behavior of allele frequency has implications for the average fitness of the population. When the mutant allele is present in frequency *x* at time *t*, the population fitness *w*(*x, t*) = (1 + *s*)*x*^2^ + 2*x*(1 − *x*)(1 + *hs*) + (1 − *x*)^2^ = 1 + *σ* sin *ωtf*(*x*) where *f*(*x*) = (1 − 2*h*)*x*^2^ + 2*hx* (see MODEL section). The population-averaged log fitness 〈ln *w*〉 oscillates about a constant which is obtained on averaging over a period of the oscillation and is given by

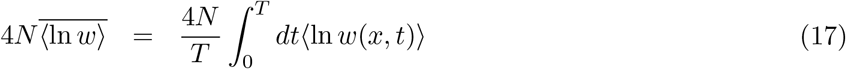

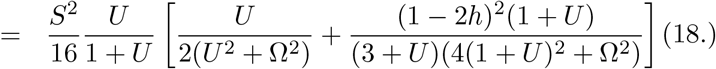

We thus find that although the selection is zero on an average, the population fitness is nonzero. The average fitness decreases with increasing driving frequency towards zero, as expected in a rapidly changing environment. We also note that the fitness is a symmetric function of *h* but the dependence is quite weak (see Fig. 5), and is apparent only at small mutation rates and for fast environmental changes.

The population fitness is also a nonmonotonic function of the scaled mutation rate *U*. For 1 ≪ *U* ≪ Ω, the fitness is close to zero because the phase difference between allele frequency and selection is large. But when *U* ≫ Ω, the phase lag decreases but the high (symmetric) mutation rate does not allow the allele frequency to deviate substantially from one half resulting in low fitness. Equation (18) also shows that the average fitness has a peak at an optimal mutation rate,

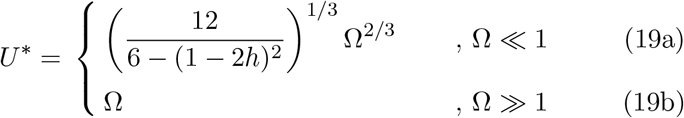

which increases with the rate of environmental change.

## 5 Discussion

In this article, we studied the evolutionary dynamics of a finite, diploid population in a varying environment for both weak and strong mutations. Assuming semi-dominance, the fixation probability of a mutant has been studied in infinitely large populations when selection changes gradually in both magnitude and direction (Uecker and Hermisson, 2011; Peischl and Kirkpartick, 2012) and in finite populations that are subjected to abrupt changes in the direction of selection (Takahata *et al*., 1975; Mustonen and Lässig, 2008; Cvijović *et al*., 2015; Dean *et al*., 2017). Here we modeled a situation in which the mutant allele is beneficial during a part of the seasonal cycle and deleterious in another, and hence its selection coefficient *s*(*t*) varies periodically with time. A key difference between our and previous body of work is that we do not assume the population to be semi-dominant.

### Rate of environmental change

Using a branching process and a diffusion theory (Uecker and Hermisson, 2011; Waxman, 2011), here we have obtained simple expressions for the fixation probability when the frequency of the environmental change is smaller or larger than the resonant frequency *ω_r_* of the population. The frequency *ω_r_* is the ‘natural frequency’ of the population and given by its growth rate when the time-averaged selection strength 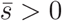 and inverse population size for 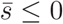. The fixation probability exhibits an extremum when the environment changes at a rate equal to the resonant frequency; whether this extremum is a minimum or a maximum depends on the arrival time *t_a_* of the mutant.

In a slowly changing environment, the fixation probability of a mutant is expected to depend on the time it arrives in the changing environment. But it is perhaps not obvious if the dependence on the initial condition remains in fast changing environments as the fixation probability in an infinitely fast changing environment is given by the corresponding result in static environment with the time-averaged selection coefficient 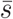. Here we find that the eventual fate of the rare mutant depends on its arrival time at any finite rate of environmental change.

Now, consider a beneficial mutant that arises when its selection strength is decreasing with time (that is, *s*(*t_a_*) > 0 but *ṡ*(*t_a_*) < 0). In a slowly changing environment, its fixation probability is smaller than that in the static environment and approaches the corresponding result in the time-averaged environment from below with increasing rate of environmental change resulting in a minimum at the resonant frequency. Thus, if the environment changes at a rate faster than the resonant frequency, a mutant that is beneficial in a static environment will have a fixation probability lower than (2*N*)^−1^ in a dynamic environment with zero or negative time-averaged selection coefficient. Similarly, a deleterious mutant in static environment can have enhanced chance of fixation in changing environments that are neutral or deleterious on an average.

The above discussion assumes that the mutations are rare. When recurrent mutations occur, we find that in the on-average neutral environment, the population can gain fitness which is, however, appreciable when the mutation rate is as high as the rate of environmental change.

### Dominance patterns in slowly changing environments

In a static environment, a dominant beneficial mutant enjoys a higher chance of fixation than a recessive one because the fitness of the mutant allele (relative to the wild type homozygote) is higher in the former case (Haldane, 1927). This pattern is reversed for deleterious mutants where the fixation of recessives is favored (Kimura, 1957). In a slowly changing environment (*ω* ≪ *ω_r_*), if the mutant starts out as a beneficial mutant (that is, its selection coefficient at arrival time *s*(*t_a_*) > 0), Haldane’s sieve operates. Similarly, if the mutant is delete-rious to begin with, its chances of fixation are reduced if it is dominant. This result is attested by Figs. 3 and S2 for 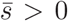, Fig. 1 for 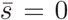 and Fig. S1 for 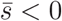.

Equations (8a) and (13b) for an on-average beneficial and neutral mutant, respectively, show that the change in fixation probability due to a slow change in the environment is simply equal to the change in the mutant’s initial fitness relative to its initial fitness, that is, *ṡ*(*t_a_*)/*s*(*t_a_*) which is *independent* of the dominance coefficient. This can be argued as follows: when the selection coefficient changes very slowly, it is reasonable to assume that the fixation probability has the same functional form as that in the static environment. Then for an initially beneficial mutant, 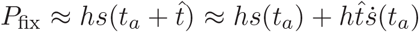 where 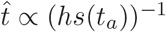 is the time at which the mutant escapes stochastic loss as estimated from a deterministic argument.

### Dominance patterns in fast changing environment

As already mentioned above, an initially beneficial mutant arising when the selection is declining can behave effectively as a deleterious mutant in a rapidly changing environment which is neutral or deleterious on an average. This has the immediate consequence that the dominant mutant is less likely to fix than the recessive one as supported by Fig. 1 for 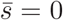 and Fig. S1 for 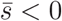. This result can be relevant to understanding adaptation in environments that change fast and for a short period of time. In Sec. S4, we construct such examples and find that an initially beneficial mutant in transiently changing environments has the same dominance patterns as discussed above in periodically changing case.

Equations (8b) and (11) show that in fast changing environments (*ω* ≫ *ω_r_*), the fixation probability is proportional to that in the infinitely fast changing environment. If genetic drift is ignored, the mutant frequency grows as *x* = *hσ* sin(*ωt*)*x* so that at large times, the average number of mutants, *n* = 1 + *hσ* cos(*θ_a_*)/*ω* for large *ω* and therefore, the fixation probability is simply that of *n* mutants (also, see Sec. S4).

### Open questions

Here we have mainly focused on the fixation probability and did not discuss how substitution rate and adaptation rate behave in changing environments. However, our preliminary simulations show that the substitution rate varies nonmonotonically with driving frequency (also see Mustonen and Lässig (2007)). A detailed understanding of these quantities requires the knowledge of fixation time which shows interesting dominance patterns in static environments (Mafessoni and Lachmann, 2015); extending such results to dynamic environment is desirable and will be discussed elsewhere.

When adaptation occurs due to standing genetic variation, the fixation probability of a beneficial mutant is known to be independent of dominance in static environments (Orr and Betancourt, 2000); here, we have studied the fixation probability of a *de novo* mutation and a detailed understanding of how standing variation affects the results obtained here is a problem for the future.

## Appendix A Branching process

The fixation probability of a mutant that arises in a large wild type population and is beneficial on an average is given by (7). For *ω* → ∞, the fixation probability is given by 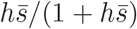 while for *ω* = 0, it is equal to *hs*(*t_a_*)/(1 + *hs*(*t_a_*)) for 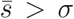 and zero otherwise (also see Fig. S2).

Away from these extreme limits, the integral appearing in (7) can be analyzed for small 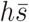, *hσ* as follows. For small frequencies 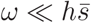, *hσ*, by first expanding the integrand in powers of *ω* and then carrying out the resulting integrals, we obtain

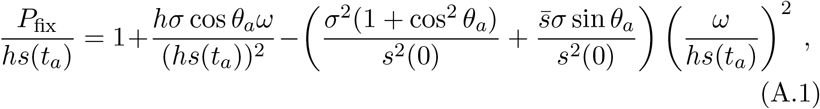

provided 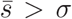 and zero otherwise. Similarly, for large frequencies, the fixation probability can be found by first expanding the integrand in powers of *hσ/ω*; for 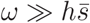, *hσ*, this finally results in

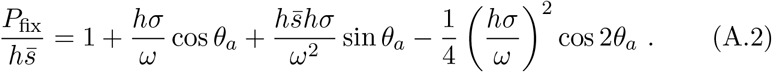

## Appendix B Fixation probability at small frequencies

Here we study (5) for 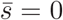 and small frequencies within a perturbation theory by writing *P*_fix_ = *P*_0_ + *NωP*_1_. It is useful to rewrite (5)

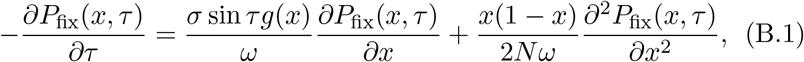

where *τ* = *ωt* + *θ_a_*, *t* ≤ 0.

In a static environment, if the mutant arises at time *t_a_* = *θ_a_*/*ω* and has fraction *x* in the population, its fixation probability is given by 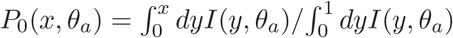, where (Kimura, 1957)

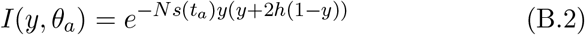

and *s*(*t_a_*) = *σ*sin*θ*_a_. For a strongly beneficial mutation (*Ns*(*t_a_*) ≫ 1), the fixation probability *P*_0_ increases with dominance coefficient and given by *hs*(*t*_*a*_), while for a deleterious mutation, it decreases with *h*. The chance of fixation also decreases with population size for *h* ≤ 1/2 but the variation with *N* is non-monotonic for *h* > 1/2.

The effect of slowly changing environment on the fixation probability is captured by *P_1_* that, by virtue of (B.1), obeys the following ordinary differential equation,

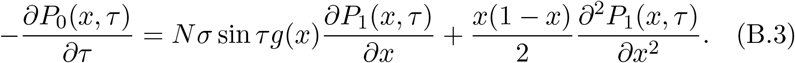

Equation (B.3) subject to boundary conditions *P*_1_(0, *τ*) = *P*_1_(1, *τ*) = 0 has the solution

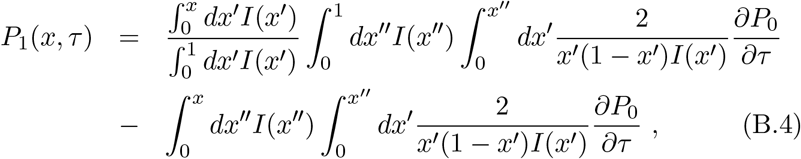

which, for small initial frequency (*τ* = *θ_a_, x* → 0) can be approximated by

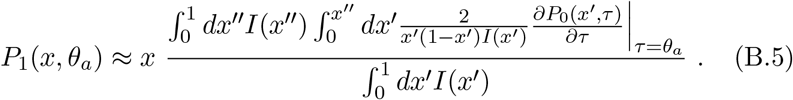

The following cases need to be considered separately:

i. −1 ≪ *Ns*(*t_a_*) ≪ 1: For small *Nσ* and arbitrary *θ_a_*, we first expand *I*(*x, τ*) to linear order in *Nσ* and carry out the integrals in the expression for *P*_0_ given above to obtain

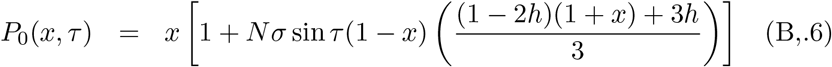

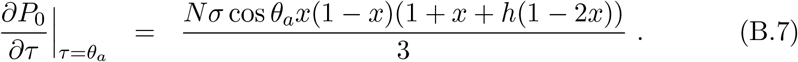 Using these approximations in (B.5), to leading order in *Nσ*, we get

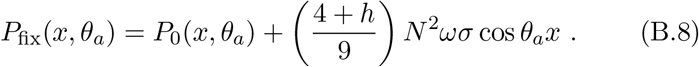 For *Ns*(*t_a_*) = 0 (that is, *θ_a_* = 0 or *π*, arbitrary *Nσ*), it can be easily seen that the function *I*(*x*, *θ_a_*) = 1 and the derivative 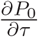 is given by (B.7) thus leading to (B.8).
ii. *Ns*(*t_a_*) ≫ 1: For large |*Ns*(*t_a_*)|, using the asymptotic expansion of the error function erf(*x*) (Abramowitz and Stegun, 1964), the fixation probability *P*_0_(*x, τ*) can be approximated as

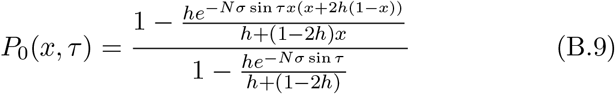

(more precisely, the above expression holds for 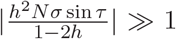). For large, positive *Ns*(*t_a_*), the denominator in (B.9) can be approximated by one leading to

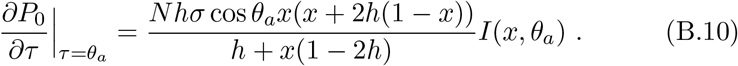 Using the above expression in (B.5) and performing the integrals for *Ns*(*t_a_*) ≫ 1, we finally obtain the following simple result,

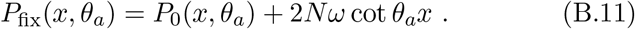
iii. *Ns*(*t_a_*) ≪ −1: Taking the derivative of *P*_0_ in (B.9) with respect to τ and keeping factors proportional to *e*^−*Nσ*sin *τ*^ only, we obtain

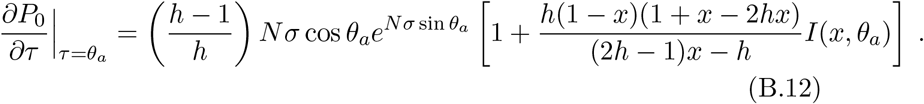 Noting that the dominant contribution to the inner integral in the numerator of (B.5) comes from *x′* → 0, we finally get

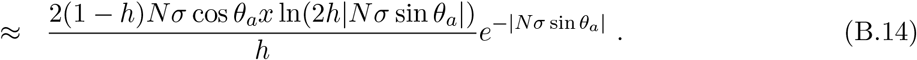

## Appendix C Fixation probability at large frequencies

Here we calculate the fixation probability of a neutral mutant 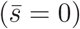 when the driving frequency is larger than the amplitude of selection (*ω* > *σ*). On writing 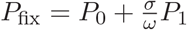 in (B.1) and collecting terms to zeroth and first order in *σ/ω*, we find that *P*_0_ = *x*, as expected. The correction *P*_1_ obeys an inhomogeneous partial differential equation,

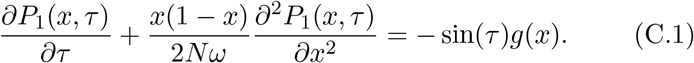

with boundary conditions *P*_1_ (0, *τ*) = *P*_1_(1, *τ*) = 0.

The homogeneous equation can be solved using standard eigenfunction expansion method (Kimura, 1955; Ewens, 2004), and we find that 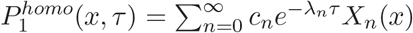 with the eigenvalue 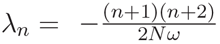 and eigenfunction 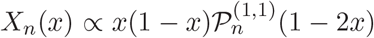 where 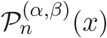 is the Jacobi polynomial (Abramowitz and Stegun, 1964). However, as this homogeneous solution is not periodic in *τ*, it does not contribute to the full solution. But since the eigenfunctions *X_n_*(*x*) form a complete set of basis, we can write 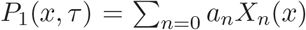 and 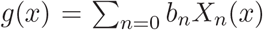 where *b_n_* are obtained using the orthogonality property of *X_n_*(*x*). Using these in (C.1), we obtain

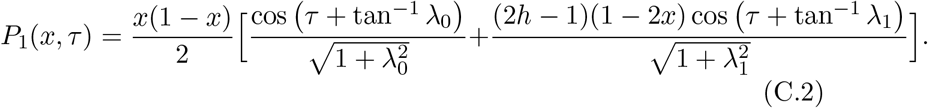

When a single mutant with frequency *x* = (2*N*)^−1^ arises at time *t_a_*, the above expression reduces to (10b) in the main text and can be used to find the resonant frequency at which the probability of fixation has an extremum. For *h* = 1/2, we find that

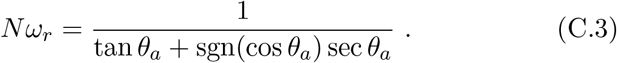

For arbitrary *h*, we are unable to find a simple closed expression for *ω_r_* as it is a solution of a 6th order algebraic equation. But a numerical study of this equation shows that *ω_r_* depends weakly on dominance coefficient.

## Appendix D Allele frequency distribution

The forward time dynamics of the population under mutation and selection are described by (14) for the allele frequency distribution Φ(*x, t*). The distribution Φ_o_ for the population subject to mutation and genetic drift is given by 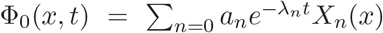 with eigenvalues 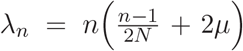 and eigenfunctions 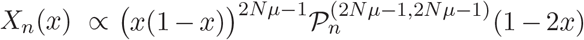 where, 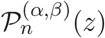 is the Jacobi polynomial (Crow and Kimura, 1956). These eigenfunctions are orthogonal with respect to the weight function *w*(*x*) = [*x*(1 — *x*)]^1–2*Nμ*^.

For weak selection (*σ* < *μ*), we can expand Φ(*x, t*) as a power series in *σ*/*μ* to write Φ = Φ_0_ + (*σ*/*μ*)Φ_1_. Using this in (14), we find that Φ_1_ obeys the following differential equation,

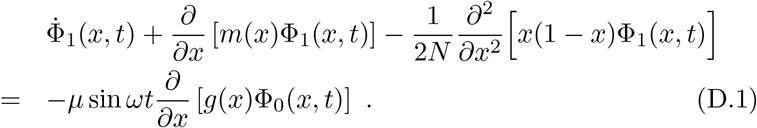

To find the distribution Φ_1_(*x, t*), we expand it and the RHS of above equation as a linear combination of *X_n_*(*x*). Writing 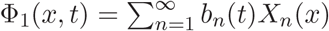 in (D.1), we find that at large times,

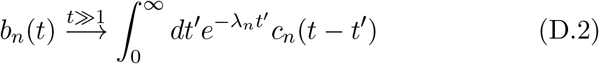

where

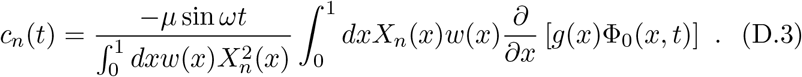

As 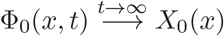, from (D.2), we obtain

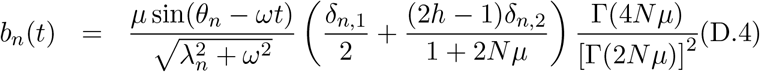

where *θ_n_* = tan ^−1^(*ω*/*λ_n_*). We thus obtain

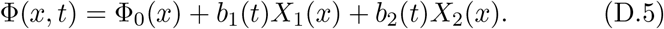

## Supporting Information

### Appendix S1 Fixation probability of a deleterious mutant

**Figure S1:**
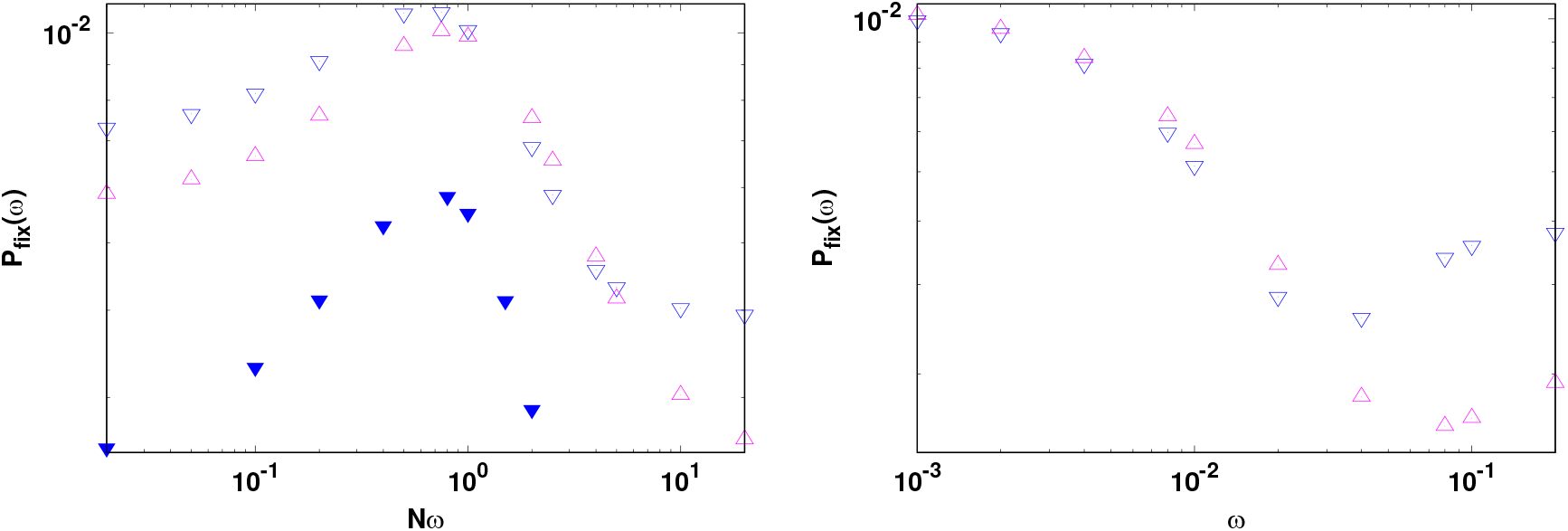
Fixation probability of a mutant which is deleterious on an average for 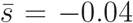, *σ* = 0.06, and *θ_a_* = *π*/8 (left panel) and 3*π*/4 (right panel). The other parameters are *N* = 50, *h* = 0.1(∇), 0.9(Δ). In the left panel, the data for *h* = 0.1, *N* = 100 (▾) is shown to support the claim that the resonant frequency scales as N^−1^.

When a mutant is deleterious at all times 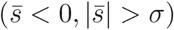, its fixation probability is lower than that of a neutral mutant in both static and dynamic environments, and the dominant mutant has a lower chance of fixation than the recessive one. But when 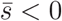 and 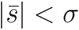, the mutant can be beneficial for some time in a periodically changing environment and its fixation probability can exceed the neutral value depending on the arrival times. In the left panel of Fig. S1, the selection coefficient *s*(*t_a_*) < 0 and therefore the recessive mutant is favored at small frequencies, while the dominant mutant has a higher chance of fixation in slowly changing environments for the parameters in the right panel since *s*(*t_a_*) > 0. In either case, at high frequencies, the fixation probability of a recessive mutant is larger since the time-averaged selection coefficient is negative. We also note that in Fig. S1a, there is a regime where the dominant mutant’s fixation probability exceeds that of the recessive one; however, the difference is quite small and a more detailed investigation is needed to evaluate the importance of this effect.

### Appendix S2 Fixation probability of a beneficial mutant

**Figure S2:**
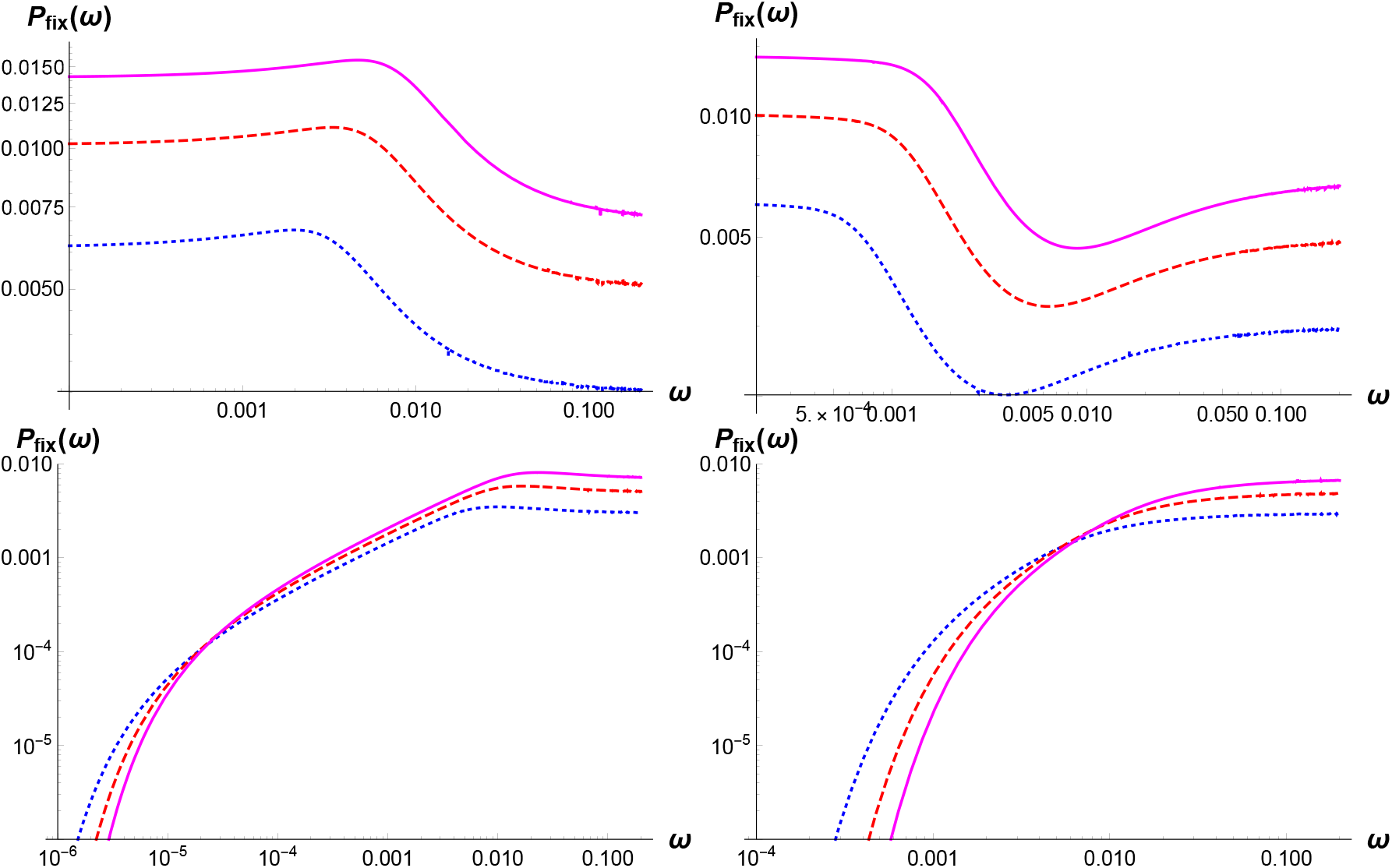
Fixation probability (7) of a mutant that is beneficial on an average for *θ_a_* = *π*/4, 3*π*/4, 5*π*/4, 7*π*/4 (clockwise from top left) for *h* = 0.3 (dotted), 0.5 (dashed), 0.7 (solid). The other parameters are 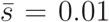 and 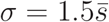.

### Appendix S3 Fixation probability of a neutral mutant

**Figure S3:**
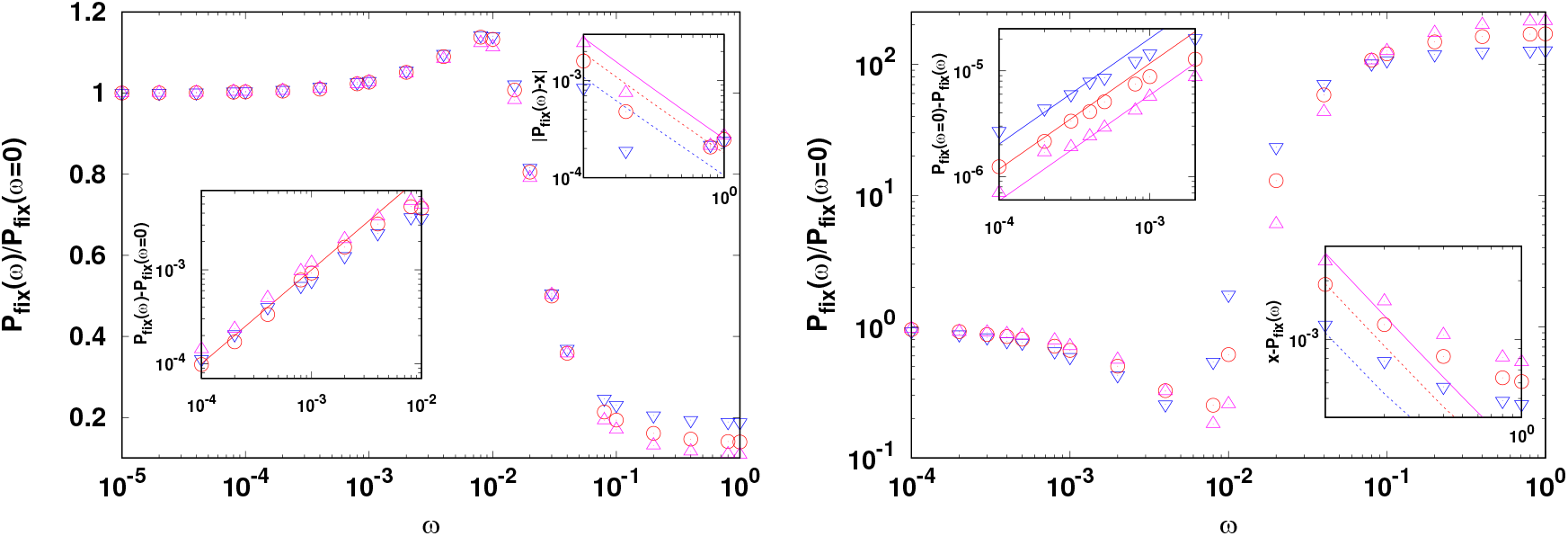
Fixation probability when the mutant is neutral on an average and *N σ* ≫ 1. The solid line shows the expression (13b) (left panel) and (13c) (right panel) for low frequencies and the dashed lines represent (10b) for high frequencies. Here *N* = 100, *σ* = 0.1, *h* = 0.3(∇), 0.5(°), 0.7(Δ), and the mutant arrived at *θ_a_* = *π*/4 (left panel) and 5*π*/4 (right panel). The numerically obtained value *P_fix_*(*ω* = 0) = 2.53 × 10^−2^, 3.40 × 10^−2^, 4.38 × 10^−2^ for *h* = 0.3, 0.5, 0.7 respectively for the left panel. For the right panel, *P*_fix_(*ω* = 0) = 3.55 × 10^−5^,2.58 × 10^−5^, 1.98 × 10^−5^ for *h* = 0.3,0.5,0.7 respectively. The simulation results are averaged over 10^7^ runs.

### Appendix S4 Transiently varying selection

**Figure S4:**
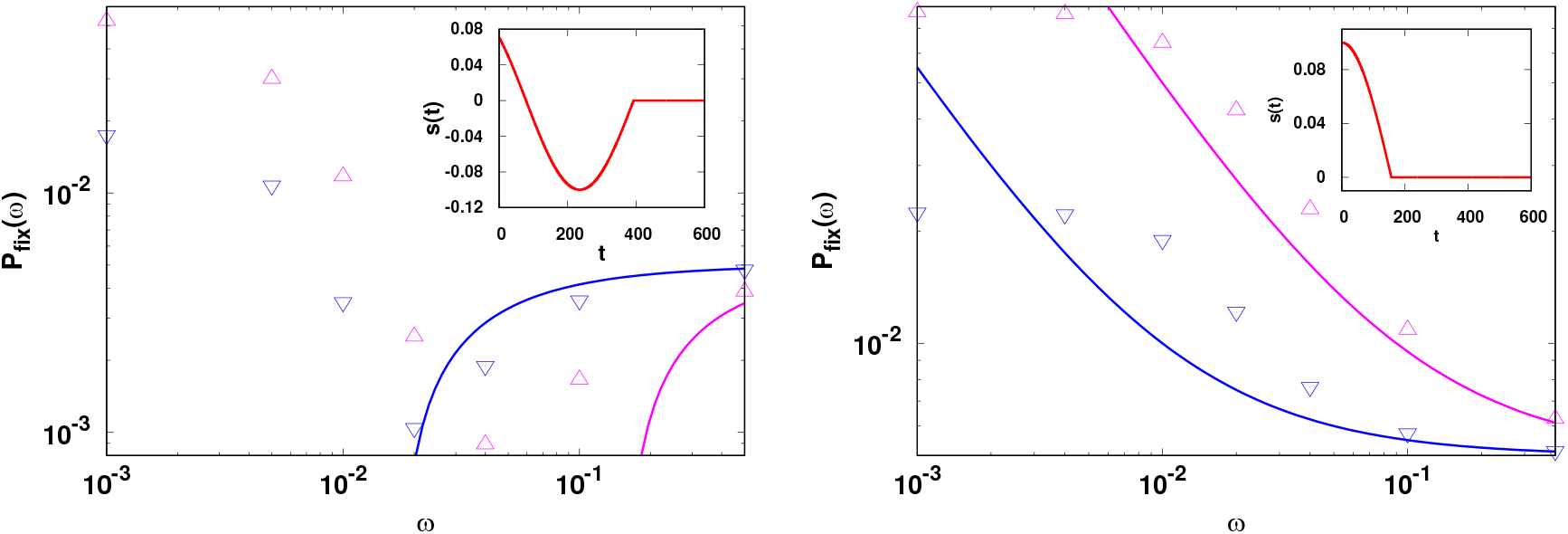
Fixation probability of a mutant in transiently varying environments defined by (S4.2) and illustrated in inset. The lines show the expression (S4.4) and points show the simulation data for *θ_a_* = 3*π*/4,*T_e_* = 2*π*/*ω* (left panel) and *θ_a_* = *π*/2,*T_e_* = *π*/*ω* (right panel). In all the plots, *N* = 100, *σ* = 0.1 and *h* = 0.1(∇), 0.9(Δ).

Here we consider a situation in which the time-averaged selection is nonzero and changes over a finite time *T_e_*,

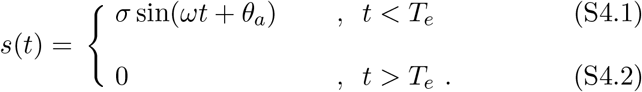

Waxman (2011) has shown that in such a case, the fixation probability is simply given by the mean allele frequency at the end of the selection. Here we estimate this allele frequency using the deterministic evolution equation,

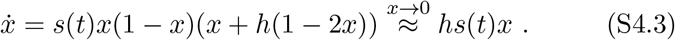

For large ω, starting from a single mutant, the number of mutants at time *T_e_* is then given by

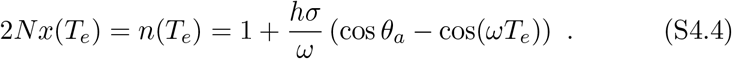

For large driving frequencies, this prediction matches qualitatively with the numerical results in Fig. S4, and therefore captures the dominance patterns when the environment changes fast over a short interval of time.

## References

Abramowitz, M. and I. A. Stegun, 1964 Handbook of Mathematical Functions with Formulas, Graphs, and Mathematical Tables. Dover.

Assaf, M., A. Kamenev, and B. Meerson, 2008 Population extinction in a time-modulated environment. Phys. Rev. E 78: 041123.

Bleuven, C. and C. R. Landry, 2016 Molecular and cellular bases of adaptation to a changing environment in microorganisms. Proc. R. Soc. B 283: 20161458. 283: 20161458.

BourGuet, D., 1999 The evolution of dominance. Heredity 83: 1–4.

Boyer, S. and G. Sherlock, 2019 Adaptation is influenced by the complexity of environmental change during evolution in a dynamic environment. bioRxiv doi:10.1101/724419-: –.

Charlesworth, B., 1998 Adaptive evolution: The struggle for dominance. Current Biology 8: R502–R504.

Crow, J. and M. Kimura, 1956 Some genetic problems in natural populations. Proc. Third Berkeley Symp. on Math. Statist. and Prob. 4: 1–22.

Cvijović, I., B. Good, E. R. Jerison, and M. M. Desai, 2015 Fate of a mutation in a fluctuating environment. Proc. Natl. Acad. Sci. USA 112: E5021–8.

Dean, A. M., C. Lehman, and X. Yi, 2017 Fluctuating selection in the Moran. Genetics 205: 1271.

Ewens, W., 2004 Mathematical Population Genetics. Springer, Berlin.

Gillespie, J. H., 1991 The Causes of Molecular Evolution. Oxford University Press, Oxford.

Gillespie, J. H., 1993 Substitution processes in molecular evolution. I. Uniform and clustered substitutions in a haploid model. Genetics 134: 971.

Haldane, J. B. S., 1927 A mathematical theory of natural and artificial selection. V. Proc. Camb. Philos. Soc. 23: 838–844.

Kimura, M., 1955 Solution of a process of random genetic drift with a continous model. Proc. Natl. Acad. Sci. USA 41: 144–150.

Kimura, M., 1957 Some Problems of Stochastic Processes in Genetics. Ann. Math. Stat. 28: 882–901.

Mafessoni, F. and M. Lachmann, 2015 Selective strolls: fixation and extinction in diploids are slower for weakly selected mutations than for neutral ones. Genetics 201: 1581–1589.

Messer, P. W., S. P. Ellner, and N. G. Hairston Jr., 2016 Can population genetics adapt to rapid evolution? Trends in Genetics 7: 408–418.

Mustonen, V. and M. Lässig, 2007 Adaptations to fluctuating selection in *Drosophila*. Proc. Natl. Acad. Sci. USA 104: 2277–2282.

Mustonen, V. and M. Lässig, 2008 Molecular evolution under fitness fluctuation. Phys. Rev. Lett. 100: 108101.

Orr, H. A. and A. J. Betancourt, 2000 Haldanes sieve and adaptation from the standing genetic variation. Genetics 157: 875–884.

Peischl, S. and M. Kirkpartick, 2012 Establishment of new mutations in changing environments. Genetics 191: 895–906.

Salignon, J., M. Richard, E. Fulcrand, H. Duplus-Bottin, and G. Yvert, 2018 Genomics of cellular proliferation in periodic environmental fluctuations. Mol Syst Biol. 14: e7823.

Takahata, N., I. Kazushige, and H. Matsuda, 1975 Effect of temporal fluctuation of selection coefficient on gene frequency in a population. Proc. Nat. Acad. Sci. USA 72: 4541.

Uecker, H. and J. Hermisson, 2011 On the fixation process of a beneficial mutation in a variable environment. Genetics 188: 915–930.

Waxman, D., 2011 A unified treatment of the probability of fixation when population size and the strength of selection change over time. Genetics 188: 907–913.

